# Alpha oscillations govern interhemispheric spike timing coordination in the honey bee brain

**DOI:** 10.1101/628867

**Authors:** Tzvetan Popov, Paul Szyszka

## Abstract

In 1929 Hans Berger discovered the alpha oscillations: a prominent, ongoing 10 Hz rhythm in the human electroencephalogram (EEG). These alpha oscillations are amongst the most widely studied cerebral signals, related to cognitive phenomena such as attention, memory, and consciousness. However, the mechanisms by which alpha oscillations affect human cognition await demonstration. Here we suggest the honey bee brain as an experimentally more accessible model system for investigating the functional roles of alpha oscillations. We found a prominent alpha wave-like spontaneous neural activity (~ 18 Hz) that is reduced in amplitude upon stimulus presentation. The phase of this alpha activity biased both timing of neuronal spikes and amplitude of high-frequency gamma activity (~ 30 Hz). These results suggest a common role of oscillatory neuronal activity across phyla and provide an unprecedented new venue for causal studies on the relationship between neuronal spikes, brain oscillations, and cognition.

## Introduction

The brain encodes information by the rate and by the timing of action potentials (spikes) across neurons. The importance of timing has already been stressed in 1929 by Hans Berger who discovered the alpha oscillations: The strongest rhythmic oscillation measurable from the human scalp EEG at a frequency of 10 Hz (Berger, 1929). In 1934 Adrian and Matthews confirmed this observation and concluded that the alpha oscillation arises “from an area in the occipital lobes connected with vision, and not from the whole cortex” (p.384) (Adrian and Matthews, 1934; Compston, 2010). Today, alpha oscillations are amongst the most widely studied psychophysiological signals. They have been linked to cognitive functions such as attention and memory in humans and other vertebrates (Klimesch et al., 2007; Palva and Palva, 2007; Jensen and Mazaheri, 2010; Jensen et al., 2014). Amplitude modulation of alpha oscillations is hypothesized to regulate neuronal excitability throughout the cortex (Klimesch et al., 2007; Jensen and Mazaheri, 2010; Hanslmayr et al., 2012), and the phase of alpha oscillations biases the rate of neuronal spiking within (Haegens et al., 2011) and across cortical areas (Saalmann et al., 2012). On a larger scale, neural processing is functionally organized in feed forward (bottom up) and feedback (top down) streams (Felleman and Van Essen, 1991). Feedback streams have been linked to alpha oscillations and feed forward streams to higher frequency oscillations above 30 Hz (gamma oscillations) (van Kerkoerle et al., 2014; Bastos et al., 2015; Michalareas et al., 2016). However, the exact mechanisms by which alpha and gamma oscillations affect neural processing and manifests in behavior still await demonstration. Progress in revealing the mechanisms and functions of oscillatory brain activity is hampered by the difficulty to relate oscillatory brain activity to the spiking activity of identified neurons in behaving animals.

In recent years insects have become model systems for studying the relationship between the spiking activity of identified neurons and the animal’s perception and cognitive performance. For example, the perceived quality of an odor can be predicted from the ensemble of co-active olfactory neurons, each being identified by its specific pattern of afferent and lateral inputs (Galizia et al., 1999; Guerrieri et al., 2005; Strutz et al., 2014; Badel et al., 2016; Grabe et al., 2016), and the mechanistic understanding of odor learning is unparalleled both in regard to molecular pathways and the identity of neuronal circuits (Hammer and Menzel, 1995; Menzel, 2012; Waddell, 2016). Similar as in vertebrates, insect brains generate oscillatory activity patterns, such as spontaneous 10 – 20 Hz oscillations in water beetles (Adrian, 1937) and honey bees (Ritz et al., 2001; Galán et al., 2004), sleep state-dependent 10 Hz oscillations in fruit flies (Yap et al., 2017), odor-induced 20 – 100 Hz oscillations in locust (Laurent and Naraghi, 1994), moth (Heinbockel et al., 1998; Daly et al., 2011) and bees (Stopfer et al., 1997; Okada and Kanzaki, 2001; Denker et al., 2010) and visual stimulus-induced 20 – 30 oscillations in fruit flies (van Swinderen and Greenspan, 2003; van Swinderen, 2007). However, albeit ubiquitous, most studies on insect brain function ignore oscillatory brain activity and focus on the encoding of sensory information via stimulus-driven changes in spike rates rather than on network-driven spike synchrony across neurons.

To investigate the temporal relationship between spikes and oscillatory brain activity within and across brain hemispheres, we performed paired recordings of local field potentials (LFP) and spikes in both mushroom bodies of the honey bee. The insect mushroom bodies are involved in olfactory learning (Erber et al., 1980; Heisenberg, 2003; Szyszka et al., 2008; Strube-Bloss et al., 2011; Menzel, 2014; Waddell, 2016), and they integrate unilaterally learned odor-reward associations across hemispheres (Strube-Bloss et al., 2016). We found an oscillatory 18 Hz activity in the LFP that exhibits properties of human alpha oscillations: 1) it was spontaneously generated, 2) it reduced amplitude upon stimulus presentation and 3) it biases the phase of both spikes and high frequency neuronal activity, effectively regulating information transmission and connectivity within and between brain hemispheres. Our data demonstrate that both spontaneous and odorant-induced spikes in the honey bee mushroom body are timed by a network-generated oscillatory clock. This oscillatory spike synchronization within and across hemispheres could play a role in object recognition as has been proposed for vertebrates (Engel et al., 1991; Mima et al., 2001) (von der Malsburg and Schneider, 1986).

## Materials and Methods

### Animals

Honey bees (*Apis mellifera*) were used in an in vivo preparation. The head capsule, thorax and abdomen were fixated with dental wax (Dr. Böhme & Schöps Dental GmbH, Deiberit 502) in a metal tube. The basis of the antennae were immobilized with eicosan (Sigma Aldrich). The cuticle of the head capsule between antennae and eyes was removed to get access to the brain. The glands in the head were removed and the head capsule was rinsed with artificial hemolymph (in mM: NaCl 130, KCl 6, Glucose 25, MgCl2 2, CaCl2 7, Sucrose 160, 10 HEPES, pH 6,7, 500 mOsmol). To avoid recording muscular activity we used Philanthotoxin-343 to paralyze the muscles in the head (50 μl, 10^-4^ Mol in artificial hemolymph; donated by P.N.R. Usherwood). Trachea between the antennal lobes and above the vertical lobes of the mushroom bodies were removed.

### Olfactory stimuli

Olfactory stimuli were applied with a computer controlled stimulator (Galizia et al., 1997). The following odorants were used: essential oils of clove, peppermint, orange (all from Apotheke Dahlem-Dorf), and geraniol, citral, isoamyl acetate und 1-heptanol (all from Sigma-Aldrich). Four μl of pure odorants were applied on 1 cm^2^ filter paper in a 1 ml syringe. The different odorants were applied alternatingly with an interstimulus interval of 10 to 15 seconds. Residual odorants were removed via an exhaust tube (5 cm inner diameter) positioned 3 cm behind the bee.

### Data acquisition

Local field potentials (LFP) and extracellular neuronal spikes were recorded in the center of the mushroom body vertical lobes (depth: 20 – 150 μm) where olfactory Kenyon cells converge onto mushroom body output neurons (Strausfeld, 2002). Recordings were performed with artificial hemolymph-filled glass microelectrodes (1/0.58 mm outer/inner diameter) that were pulled with a micropipette puller (P-2000, Sutter Instrument CO) to get a tip resistance of 5 to 10 MOhm and which were then broken to get a tip resistance of 1.3 – 3.5 MOhm. Spikes could only be recorded with electrodes that had a tip resistance between 2.4 to 3.5 MOhm (which was the case in 5 out of the 10 recorded bees). The reference electrode was a chloridized silver wire (0.2 mm diameter). The recording chain consisted of a 10x-amplifier (Simmonds-Amplifier, Cambridge, UK), a 1000x-amplifier and 0.1 – 3.000 Hz band-pass filter (AM503, Tektronix) and an analog-digital converter (1401+, Science Park Cambridge, UK). Experiments were performed at room temperature.

### Data analysis

Data analysis was performed with the MATLAB FieldTrip toolbox (Oostenveld et al., 2011). Spectral analysis was computed for each trial using a Fast Fourier Transformation (FFT) utilizing a multi-taper approach with orthogonal Slepian tapers (Mitra and Pesaran, 1999). For frequencies below 40 Hz, 3 orthogonal Slepian tapers were used, resulting in frequency smoothing of ±2 Hz. Spectrally resolved Granger causality (GC) analysis (Granger, 1969; Ding et al., 2006; Wen et al., 2013) was used to identify the directionality (so-called feedforward vs. feedback influences) of information flow between cortical areas (Figure 2C). The first .5 s post stimulus onset was omitted in order to avoid transient responses biasing Granger estimates. A bivariate nonparametric spectral matrix factorization approach (Wen et al., 2013) was applied in order to estimate GC. This algorithm estimates the spectral density matrix on the basis of Fourier coefficients that were computed using multitapers as described above (3 orthogonal Slepian tapers) with frequency smoothening of ±3 Hz for frequencies of 0 to 100 Hz with 1 Hz spectral resolution. On the basis of the spectral density matrix (i.e., applying spectral factorization), the noise covariance matrix and the transfer function were obtained (Wen et al., 2013).

Neural spikes were identified from the raw LFP recordings using the peak detection algorithm implemented in MATLAB -*findpeaks.m*. Spikes were identified as local maxima exceeding 0.1 mV as illustrated in Figure S2. Subsequently the indices of these local maxima were used as triggers in order to re-segment the data ± 1sec around each spike (Figure 2A, B).

Spike rate as a function of the spontaneous oscillatory phase was determined as follows (Figure 2E). First, the individual peak frequency in the range 10-25 Hz was identified and the analytic amplitude was computed on the band pass (±1 Hz around the individual peak frequency) filtered epoched data. This step allows the derivation of phase information per time sample. Data intervals corresponding to odor presentation (−.2 to 2.2 sec) and spontaneous activity (−4 to 0 sec) were re-segmented. Subsequently, these segments were vertically concatenated. For the filtered data, concatenation was performed after multiplication with a Hanning window to minimize the effect of edge artifacts at the beginning and the end of the initially odor onset epoched data. Finally, the indices of spikes were identified as described above together with their corresponding phase value. Effects of spontaneous oscillatory power on spike rate were evaluated by comparing high vs. low power data segments in a Bee × Latency (spontaneous vs. odor stimulation or high vs. low spontaneous power) analysis of variance (ANOVA). Significant main effects and interactions were decomposed by simple effects ANOVA or *t* test.

## Results

LFP recordings in the vertical lobes of the honey bee mushroom bodies revealed a spontaneous ~18 Hz oscillation in both hemispheres (Figure 1A). The amplitude of the 18 Hz oscillatory activity was reduced upon odor stimulation and returned back to baseline shortly after (Figure 1B). This power reduction was accompanied by a stimulus-induced power increase in 20-40 Hz band (low gamma oscillation) (Figure 1C). In addition, there was an increase of power at frequencies above 40 Hz (high gamma activity) (Figure 1D). This high gamma activity coincided with the stimulus-induced low gamma oscillation. Increases in high gamma power are thought to reflect the increase in the underlying spiking activity of multiple neurons (Ray et al., 2008; Ray and Maunsell, 2011). Both, the spontaneous alpha oscillation and the odor-induced low and high gamma activity required an intact antenna ipsilateral to the recording site (Figure S1). This suggests that both alpha and gamma oscillations are generated in or require input from the ipsilateral antennal lobe (first olfactory brain center and functional analog of the vertebrate olfactory bulb).

**Figure 1:**
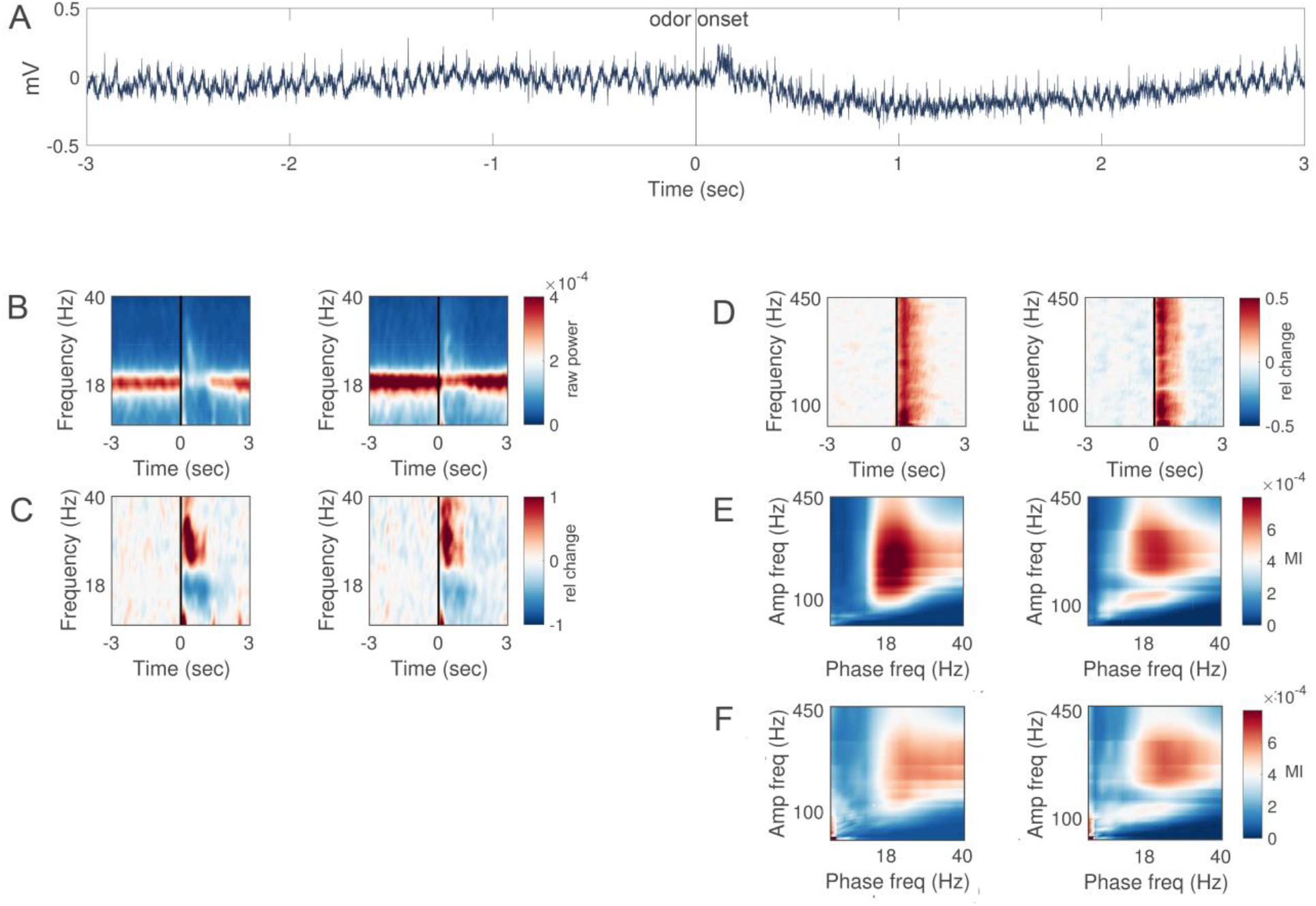
Spontaneous alpha and odor-induced gamma oscillations in the mushroom bodies. **A:** Raw trace of a single trial in a representative bee. Odor is presented at 0 sec. **B**: Time-frequency (5-40Hz, alpha and low gamma) representation of raw power (color-coded, mV^2^/Hz) recorded in the left (left) and right hemisphere (right) respectively. An odor stimulus was presented at 0 sec for the duration of 1 sec. N = 10 bees (stimulations per bee mean/SD 52.6/28.9). **C:** Same as in B baseline corrected. Warm colors indicate increase and cold colors decrease in oscillatory power expressed as relative change from pre-stimulus baseline. **D**: Same as in C but for higher frequencies (40-450Hz, high gamma). **E**: Cross-frequency phase-to-amplitude coupling. Bispectra illustrating cross-frequency relationships in the left and right hemisphere respectively. The phase providing frequency is depicted on the x-axis and amplitude providing frequency on the y-axis. Color code depict the modulation index (MI). **F**: Same as E but during olfactory stimulation.

The cross-frequency coupling of high gamma activity to the phase of the spontaneous alpha oscillation is considered a mechanism of neural communication in cortical circuits (Canolty and Knight, 2010). Analysis of the cross-frequency coupling between high gamma activity and alpha oscillations confirms that the high gamma amplitude is modulated by the phase of the spontaneous alpha oscillation around 18 Hz (Figure 1E). Moreover, there was also a cross-frequency coupling between high gamma activity and odor-induced gamma oscillations (Figure 1F).

In human EEG recordings such cross-frequency coupling between high gamma activity and alpha oscillations is typically interpreted as an indicator of stimulus- or task-induced changes in neuronal spiking that is biased by the phase of the spontaneous alpha rhythm (Haegens et al., 2011). Unlike in human EEG, in honey bees the accesses to both, LFP and neuronal spikes allow the empirical test of this hypothesis. An affirmative confirmation of the coupling between spikes and alpha oscillations is illustrated by the spike-field coherence analysis in Figure 2. A reliable 18 Hz peak in the coherence spectrum was visible in both hemispheres, suggesting that the timing of spikes is coupled to the phase of the spontaneous 18 Hz oscillation. The timing of spikes in the right hemisphere was phase-coupled to the 18 Hz oscillation in the left hemisphere (Figure 2A). In contrast, the timing of spikes in the left hemisphere was not phase-coupled to the 18 Hz oscillation in the right hemisphere (Figure 2B). In other words, the direction of information flow quantified by spike transmission was stronger in the direction from the right to left hemisphere. This right-over-left dominance in interhemispheric information flow was also visible in an analysis of Granger causality (Granger, 1969; Wen et al., 2013) (Figure 2C).

**Figure 2:**
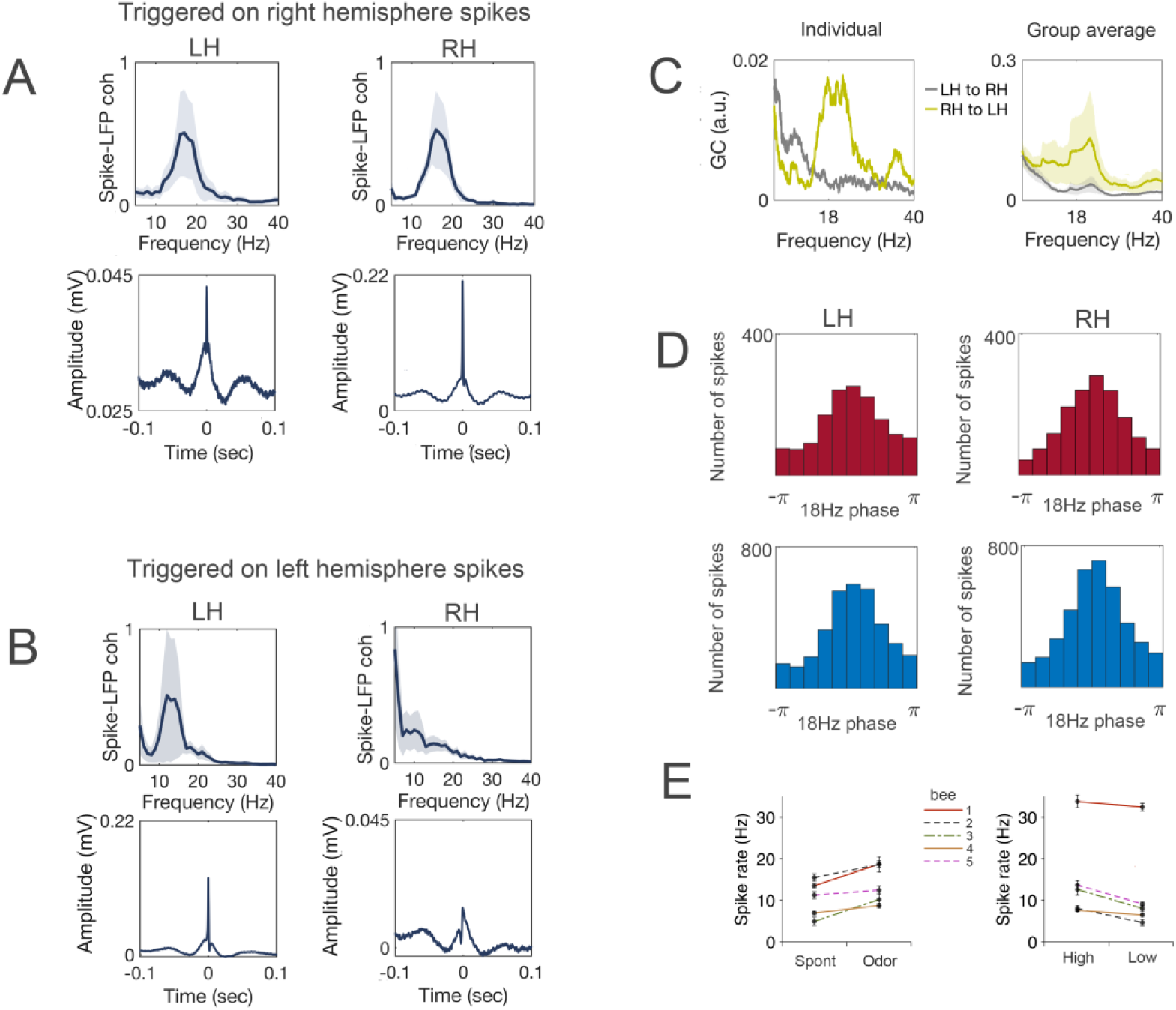
Neuronal spikes are phase-locked to the alpha oscillation. **A:** Spike-triggered averages computed on right hemisphere spikes. Top-Coherence between spikes and low gamma activity computed over the spike triggered averages indicating a reliable ~18 Hz peak in the spectrum (mean and SEM, N = 5 bees). Bottom-Spike-triggered averages for the left and right hemisphere respectively. Spike onset is evident at 0 sec. N = 5 bees (spikes per bee mean/SD 4441/2415). **B:** Same as A computed on spikes identified in the left hemisphere. **C:** Granger causality spectra for an individual bee (left) and the group average indicates stronger Granger causality influence from the right hemisphere (RH) to the left hemisphere (LH) (N = 10 bees, line represents the mean, the shaded area the SEM, right). **D:** Phase preference of neuronal spikes. Distribution of the spikes within the 18 Hz oscillatory cycle. Top-spikes identified during the odor stimulation interval. Bottom-spikes identified during the pre-stimulus baseline period. X-axis denotes the phase and y-axis the absolute number of identified spikes. **E:** Left: Spike rate per bee during spontaneous activity (during 2 to 1 sec before odor onset) and odor evoked activity (during 0.1 to 1.1 sec after odor onset). Right: Spike rate per bee for high and low alpha power. Points and error bars show the mean and SED over trials. Values show number of spikes per second.

Neuronal spikes were biased by the 18 Hz phase and displayed a non-uniform phase preference, and the recurrence of spikes was larger during the peak of the alpha oscillation during both spontaneous and odor-induced activity (Figure 2D). During spontaneous activity the spike rate depended on the power of the alpha oscillation, and was higher in epochs of high alpha power (Figure 2E right) as confirmed by the main effect Latency (F(1,341) = 18.74, p < 0.001). There was a main effect for Bee (F(4,341) = 32.27, p < 0.001) and no Bee × Latency interaction. Moreover, the spike rate was higher during odor-induced activity than during spontaneous activity (Figure 2E left, F(1,341) = 32.05, p < 0.001), corresponding to the period characterized by the strongest decrease in alpha power (Figure 1B and C).

## Discussion

Here we demonstrate that the LFP activity in the brain of the honey bee shares characteristics of EEG activity in the human brain. First, there is a spontaneously generated, ongoing alpha oscillation with a characteristic peak in the spectrum (18 Hz in bees, 10 Hz in humans). Second, the amplitude of the alpha oscillation is reduced upon stimulus presentation. Third, the phase of the spontaneous alpha oscillation biases both high-gamma oscillatory activity and neuronal spiking. Fourth, the spike rate depends on the power of the alpha oscillation. However, opposite to vertebrates (Haegens et al., 2011), in bees the spike rates decrease with a decrease in α-power.

We recorded LFPs and spikes in the output region of the mushroom bodies (vertical lobes), where several thousand mushroom body-intrinsic Kenyon cells converge onto a few hundred output neurons (Rybak and Menzel, 1993; Strausfeld, 2002). LFPs in the vertical lobes are likely generated by the summation of excitatory postsynaptic potentials across mushroom body output neurons (Kaulen et al., 1984). However, it is not possible to determine the number or the identity of output neurons that contributed to the LFP oscillations and spikes. Previous studies showed that odor-evoked 30 Hz oscillations in the mushroom body input region (calyx) are driven by oscillatory spike synchronization across projection neurons which connect the antennal lobe with the ipsilateral mushroom body (Laurent and Davidowitz, 1994; Stopfer et al., 1997). The fact that the odor-evoked 30 Hz oscillations, which we recorded in the mushroom body output region, require input from the ipsilateral antenna, suggests that they also reflect oscillatory spike synchronization across projection neurons. Likewise, the spontaneous 18 Hz oscillation required an intact ipsilateral antenna, indicating that it is also driven by projection neurons of the ipsilateral antennal lobe. This hypothesis is supported by the fact that projection neurons are spontaneously active (Galan et al., 2006), and this spontaneous activity is driven by antennal input and is reduced upon odor stimulation (Paoli et al., 2016). The coupling of spontaneous oscillations across hemispheres suggests, that they are generated by circuits that involve bilateral neurons which connect both antennal lobes (Abel et al., 2001) or mushroom bodies (Rybak and Menzel, 1993; Strausfeld, 2002).

The effective connectivity between cortical modules (left and right mushroom bodies) could be established based on both spike occurrence and Granger causality analyses which indicate a right-over-left dominance in interhemispheric information flow during spontaneous activity. This right-over-left dominance in interhemispheric information flow could be related to honey bees’ left-right-asymmetry of odor-reward learning (Letzkus et al., 2006) and odor discrimination at the behavioral and neuronal level (Rigosi et al., 2015). The interhemispheric coordination of spike timing could serve to bind bilateral olfactory information into coherent object representations (von der Malsburg and Schneider, 1986) and could for example underlie bees’ ability to retrieve unilaterally learned odor-reward associations via both antennae (Sandoz and Menzel, 2001; Strube-Bloss et al., 2016). Given that honey bees show cognitive capacities (e.g., concept learning (Giurfa et al., 2001); selective attention (Paulk et al., 2014); map-like spatial memories (Menzel et al., 2005)), our results suggest the honey bee as animal model to examine the functional role of oscillatory brain activity in perception and cognitive function. The ease with which brain oscillations and spikes can be recorded in honey bees opens opportunities inaccessible in human electrophysiology.

## Acknowledgments

We thank Randolf Menzel for discussions initiating the experiments and for his support during data acquisition. We thank Peter N. R. Usherwood for donating Philanthotoxin-343. This manuscript has been released as a Pre-Print at https://www.biorxiv.org/ (Popov and Szyszka, 2019).

## Author contributions

PS performed the experiments. TP analyzed the data. TP and PS wrote the manuscript.

## Supplementary Figures

**Figure S 1:**
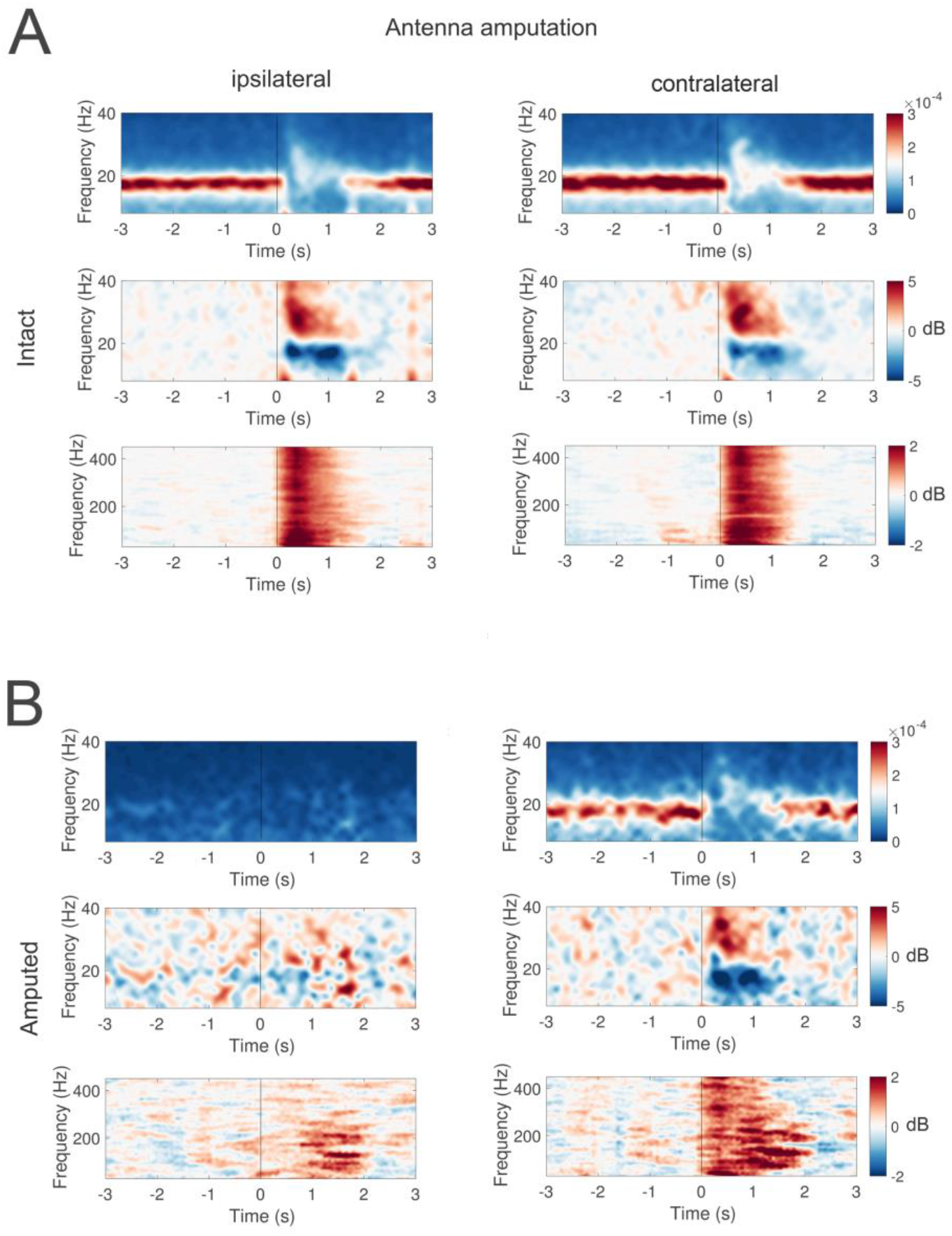
Spontaneous alpha and stimulus-induced gamma oscillations in the mushroom body require input from the ipsilateral antennal lobe. **A:** time-frequency representation of raw power below 40 Hz (alpha and low gamma) (top), baseline corrected power below (40 Hz) (middle) and baseline corrected power above 40 Hz (high gamma) (bottom) in the mushroom body ipsilateral (left) and contralateral (right) to the side of antenna amputation. N = 5 bees (stimulations per bee mean/SD 23.2/17.9 **B**: same as in A but after antenna amputation. Left: ipsilateral to antenna amputation; right: contralateral to antenna amputation. N = 5 bees (stimulations per bee mean/SD 8.4/3.4).

**Figure S 2:**
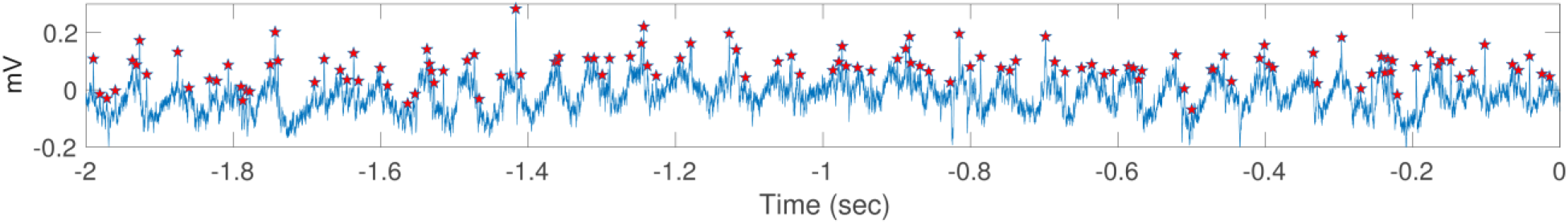
Spike detection. Raw LFP trace of the pre-stimulus baseline corresponding to the same trial illustrated in Fig 1. Red stars denote the identified spikes.

## References

Abel, R., Rybak, J., and Menzel, R. (2001). Structure and response patterns of olfactory interneurons in the honeybee, Apis mellifera. J Comp Neurol 437(3), 363–383.

Adrian, E.D. (1937). Synchronized reactions in the optic ganglion of dytiscus. J Physiol 91(1), 66–89.

Adrian, E.D., and Matthews, B.H.C. (1934). The Berger rythm: potential changes from the occipital lobes in man. Brain 57(4), 355–385. doi: https://doi.org/10.1093/brain/57.4.355.

Badel, L., Ohta, K., Tsuchimoto, Y., and Kazama, H. (2016). Decoding of Context-Dependent Olfactory Behavior in Drosophila. Neuron 91(1), 155–167. doi: 10.1016/j.neuron.2016.05.022.

Bastos, A.M., Vezoli, J., Bosman, C.A., Schoffelen, J.M., Oostenveld, R., Dowdall, J.R., et al. (2015). Visual areas exert feedforward and feedback influences through distinct frequency channels. Neuron 85(2), 390–401. doi: 10.1016/j.neuron.2014.12.018.

Berger, H. (1929). Über das Elektrenkephalogramm des Menschen. . Archiv f. Psychiatrie 87, 527. doi: https://doi.org/10.1007/BF01797193.

Canolty, R.T., and Knight, R.T. (2010). The functional role of cross-frequency coupling. Trends Cogn Sci 14(11), 506–515. doi: 10.1016/j.tics.2010.09.001.

Compston, A. (2010). The Berger rhythm: potential changes from the occipital lobes in man. Brain 133(Pt 1), 3–6.

Daly, K.C., Galan, R.F., Peters, O.J., and Staudacher, E.M. (2011). Detailed Characterization of Local Field Potential Oscillations and Their Relationship to Spike Timing in the Antennal Lobe of the Moth Manduca sexta. Front Neuroeng 4, 12. doi: 10.3389/fneng.2011.00012.

Denker, M., Finke, R., Schaupp, F., Grun, S., and Menzel, R. (2010). Neural correlates of odor learning in the honeybee antennal lobe. Eur J Neurosci 31(1), 119–133.

Ding, M., Chen, Y., and Bressler, S. (2006). “Granger Causality: Basic Theory and Application to Neuroscience.,” in Handbook of Time Series Analysis,, eds. B. Schelter, M. Winterhalder & J. Timmer. Wiley-VCH Verlage), 451–474.

Engel, A.K., Konig, P., Kreiter, A.K., and Singer, W. (1991). Interhemispheric synchronization of oscillatory neuronal responses in cat visual cortex. Science 252(5009), 1177–1179.

Erber, J., Masuhr, T., and Menzel, R. (1980). Localization of short-term memory in the brain of the bee, Apis mellifera. Physiological Entomology 5(4). doi: https://doi.org/10.1111/j.1365-3032.1980.tb00244.x.

Felleman, D.J., and Van Essen, D.C. (1991). Distributed hierarchical processing in the primate cerebral cortex. Cereb Cortex 1(1), 1–47.

Galán, R., Ritz, R., Herz, A., and Szyszka, P. (2004). Uncovering Short-Time Correlations Between Multichannel Recordings of Brain Activity: a Phase-Space Approach. Int. J. Bifurc. Chaos 14(02), 585–597.

Galan, R.F., Weidert, M., Menzel, R., Herz, A.V., and Galizia, C.G. (2006). Sensory memory for odors is encoded in spontaneous correlated activity between olfactory glomeruli. Neural Comput 18(1), 10–25. doi: 10.1162/089976606774841558.

Galizia, C.G., Joerges, J., Kuttner, A., Faber, T., and Menzel, R. (1997). A semi-in-vivo preparation for optical recording of the insect brain. J Neurosci Methods 76(1), 61–69.

Galizia, C.G., Sachse, S., Rappert, A., and Menzel, R. (1999). The glomerular code for odor representation is species specific in the honeybee Apis mellifera. Nat Neurosci 2(5), 473–478. doi: 10.1038/8144.

Giurfa, M., Zhang, S., Jenett, A., Menzel, R., and Srinivasan, M.V. (2001). The concepts of ‘sameness’ and ‘difference’ in an insect. Nature 410(6831), 930–933. doi: 10.1038/35073582.

Grabe, V., Baschwitz, A., Dweck, H.K.M., Lavista-Llanos, S., Hansson, B.S., and Sachse, S. (2016). Elucidating the Neuronal Architecture of Olfactory Glomeruli in the Drosophila Antennal Lobe. Cell Rep 16(12), 3401–3413. doi: 10.1016/j.celrep.2016.08.063.

Granger, C.W.J. (1969). Investigating Causal Relations by Econometric Models and Cross-spectral Methods. Econometrica 37(3), 424–438.

Guerrieri, F., Schubert, M., Sandoz, J.C., and Giurfa, M. (2005). Perceptual and neural olfactory similarity in honeybees. PLoS Biol 3(4), e60. doi: 10.1371/journal.pbio.0030060.

Haegens, S., Nacher, V., Luna, R., Romo, R., and Jensen, O. (2011). alpha-Oscillations in the monkey sensorimotor network influence discrimination performance by rhythmical inhibition of neuronal spiking. Proc Natl Acad Sci U S A 108(48), 19377–19382. doi: 10.1073/pnas.1117190108.

Hammer, M., and Menzel, R. (1995). Learning and memory in the honeybee. J Neurosci 15(3 Pt 1), 1617–1630.

Hanslmayr, S., Staudigl, T., and Fellner, M.C. (2012). Oscillatory power decreases and long-term memory: the information via desynchronization hypothesis. Front Hum Neurosci 6, 74. doi: 10.3389/fnhum.2012.00074.

Heinbockel, T., Kloppenburg, P., and Hildebrand, J.G. (1998). Pheromone-evoked potentials and oscillations in the antennal lobes of the sphinx moth Manduca sexta. J Comp Physiol A 182(6), 703–714.

Heisenberg, M. (2003). Mushroom body memoir: from maps to models. Nat Rev Neurosci 4(4), 266–275. doi: 10.1038/nrn1074.

Jensen, O., Gips, B., Bergmann, T.O., and Bonnefond, M. (2014). Temporal coding organized by coupled alpha and gamma oscillations prioritize visual processing. Trends Neurosci 37(7), 357–369. doi: 10.1016/j.tins.2014.04.001.

Jensen, O., and Mazaheri, A. (2010). Shaping functional architecture by oscillatory alpha activity: gating by inhibition. Front Hum Neurosci 4, 186. doi: 10.3389/fnhum.2010.00186.

Kaulen, P., Erber, J., and Mobbs, P. (1984). Current source-density analysis in the mushroom bodies of the honeybee (Apis mellifera carnica). Journal of Comparative Physiology A 154(4), 569–582.

Klimesch, W., Sauseng, P., and Hanslmayr, S. (2007). EEG alpha oscillations: the inhibition-timing hypothesis. Brain Res Rev 53(1), 63–88. doi: 10.1016/j.brainresrev.2006.06.003.

Laurent, G., and Davidowitz, H. (1994). Encoding of olfactory information with oscillating neural assemblies. Science 265(5180), 1872–1875. doi: 10.1126/science.265.5180.1872.

Laurent, G., and Naraghi, M. (1994). Odorant-induced oscillations in the mushroom bodies of the locust. J Neurosci 14(5 Pt 2), 2993–3004.

Letzkus, P., Ribi, W.A., Wood, J.T., Zhu, H., Zhang, S.W., and Srinivasan, M.V. (2006). Lateralization of olfaction in the honeybee Apis mellifera. Curr Biol 16(14), 1471–1476. doi: 10.1016/j.cub.2006.05.060.

Menzel, R. (2012). The honeybee as a model for understanding the basis of cognition. Nat Rev Neurosci 13(11), 758–768. doi: 10.1038/nrn3357.

Menzel, R. (2014). The insect mushroom body, an experience-dependent recoding device. J Physiol Paris 108(2–3), 84–95. doi: 10.1016/j.jphysparis.2014.07.004.

Menzel, R., Greggers, U., Smith, A., Berger, S., Brandt, R., Brunke, S., et al. (2005). Honey bees navigate according to a map-like spatial memory. Proc Natl Acad Sci U S A 102(8), 3040–3045. doi: 10.1073/pnas.0408550102.

Michalareas, G., Vezoli, J., van Pelt, S., Schoffelen, J.M., Kennedy, H., and Fries, P. (2016). Alpha-Beta and Gamma Rhythms Subserve Feedback and Feedforward Influences among Human Visual Cortical Areas. Neuron 89(2), 384–397. doi: 10.1016/j.neuron.2015.12.018.

Mima, T., Oluwatimilehin, T., Hiraoka, T., and Hallett, M. (2001). Transient interhemispheric neuronal synchrony correlates with object recognition. J Neurosci 21(11), 3942–3948.

Mitra, P.P., and Pesaran, B. (1999). Analysis of dynamic brain imaging data. Biophys J 76(2), 691–708. doi: 10.1016/S0006-3495(99)77236-X.

Okada, K., and Kanzaki, R. (2001). Localization of odor-induced oscillations in the bumblebee antennal lobe. Neurosci Lett 316(3), 133–136. doi: 10.1016/s0304-3940(01)02385-0.

Oostenveld, R., Fries, P., Maris, E., and Schoffelen, J.M. (2011). FieldTrip: Open source software for advanced analysis of MEG, EEG, and invasive electrophysiological data. Comput Intell Neurosci 2011, 156869. doi: 10.1155/2011/156869.

Palva, S., and Palva, J.M. (2007). New vistas for alpha-frequency band oscillations. Trends Neurosci 30(4), 150–158. doi: 10.1016/j.tins.2007.02.001.

Paoli, M., Weisz, N., Antolini, R., and Haase, A. (2016). Spatially resolved time-frequency analysis of odour coding in the insect antennal lobe. Eur J Neurosci 44(6), 2387–2395. doi: 10.1111/ejn.13344.

Paulk, A.C., Stacey, J.A., Pearson, T.W., Taylor, G.J., Moore, R.J., Srinivasan, M.V., et al. (2014). Selective attention in the honeybee optic lobes precedes behavioral choices. Proc Natl Acad Sci U S A 111(13), 5006–5011. doi: 10.1073/pnas.1323297111.

Popov, T., and Szyszka, P. (2019). Alpha oscillations govern interhemispheric spike timing coordination in the honey bee brain. BioRxiv 628867. doi: https://doi.org/10.1101/628867.

Ray, S., Crone, N.E., Niebur, E., Franaszczuk, P.J., and Hsiao, S.S. (2008). Neural correlates of high-gamma oscillations (60-200 Hz) in macaque local field potentials and their potential implications in electrocorticography. J Neurosci 28(45), 11526–11536. doi: 10.1523/JNEUROSCI.2848-08.2008.

Ray, S., and Maunsell, J.H. (2011). Different origins of gamma rhythm and high-gamma activity in macaque visual cortex. PLoS Biol 9(4), e1000610. doi: 10.1371/journal.pbio.1000610.

Rigosi, E., Haase, A., Rath, L., Anfora, G., Vallortigara, G., and Szyszka, P. (2015). Asymmetric neural coding revealed by in vivo calcium imaging in the honey bee brain. Proc Biol Sci 282(1803), 20142571. doi: 10.1098/rspb.2014.2571.

Ritz, R., Galán, R., Szyszka, P., and Herz, A. (2001). Analysis of odor processing in the mushroom bodies of the honeybee. Neurocomputing 38–40, 313–318.

Rybak, J., and Menzel, R. (1993). Anatomy of the mushroom bodies in the honey bee brain: the neuronal connections of the alpha-lobe. J Comp Neurol 334(3), 444–465. doi: 10.1002/cne.903340309.

Saalmann, Y.B., Pinsk, M.A., Wang, L., Li, X., and Kastner, S. (2012). The pulvinar regulates information transmission between cortical areas based on attention demands. Science 337(6095), 753–756. doi: 10.1126/science.1223082.

Sandoz, J.C., and Menzel, R. (2001). Side-specificity of olfactory learning in the honeybee: generalization between odors and sides. Learn Mem 8(5), 286–294. doi: 10.1101/lm.41401.

Stopfer, M., Bhagavan, S., Smith, B.H., and Laurent, G. (1997). Impaired odour discrimination on desynchronization of odour-encoding neural assemblies. Nature 390(6655), 70–74. doi: 10.1038/36335.

Strausfeld, N.J. (2002). Organization of the honey bee mushroom body: representation of the calyx within the vertical and gamma lobes. J Comp Neurol 450(1), 4–33. doi: 10.1002/cne.10285.

Strube-Bloss, M.F., Nawrot, M.P., and Menzel, R. (2011). Mushroom body output neurons encode odor-reward associations. J Neurosci 31(8), 3129–3140. doi: 10.1523/JNEUROSCI.2583-10.2011.

Strube-Bloss, M.F., Nawrot, M.P., and Menzel, R. (2016). Neural correlates of side-specific odour memory in mushroom body output neurons. Proc Biol Sci 283(1844). doi: 10.1098/rspb.2016.1270.

Strutz, A., Soelter, J., Baschwitz, A., Farhan, A., Grabe, V., Rybak, J., et al. (2014). Decoding odor quality and intensity in the Drosophila brain. Elife 3, e04147. doi: 10.7554/eLife.04147.

Szyszka, P., Galkin, A., and Menzel, R. (2008). Associative and non-associative plasticity in kenyon cells of the honeybee mushroom body. Front Syst Neurosci 2, 3. doi: 10.3389/neuro.06.003.2008.

van Kerkoerle, T., Self, M.W., Dagnino, B., Gariel-Mathis, M.A., Poort, J., van der Togt, C., et al. (2014). Alpha and gamma oscillations characterize feedback and feedforward processing in monkey visual cortex. Proc Natl Acad Sci U S A 111(40), 14332–14341. doi: 10.1073/pnas.1402773111.

van Swinderen, B. (2007). Attention-like processes in Drosophila require short-term memory genes. Science 315(5818), 1590–1593. doi: 10.1126/science.1137931.

van Swinderen, B., and Greenspan, R.J. (2003). Salience modulates 20-30 Hz brain activity in Drosophila. Nat Neurosci 6(6), 579–586. doi: 10.1038/nn1054.

von der Malsburg, C., and Schneider, W. (1986). A neural cocktail-party processor. Biol Cybern 54(1), 29–40.

Waddell, S. (2016). Neural Plasticity: Dopamine Tunes the Mushroom Body Output Network. Curr Biol 26(3), R109–112. doi: 10.1016/j.cub.2015.12.023.

Wen, X., Rangarajan, G., and Ding, M. (2013). Multivariate Granger causality: an estimation framework based on factorization of the spectral density matrix. Philos Trans A Math Phys Eng Sci 371(1997), 20110610. doi: 10.1098/rsta.2011.0610.

Yap, M.H.W., Grabowska, M.J., Rohrscheib, C., Jeans, R., Troup, M., Paulk, A.C., et al. (2017). Oscillatory brain activity in spontaneous and induced sleep stages in flies. Nat Commun 8(1), 1815. doi: 10.1038/s41467-017-02024-y.

